# FOXI3 Promotes Migration and Proliferation in Prostate Cancer Bone Metastases, Modulated by FGF8

**DOI:** 10.1101/2025.10.22.683826

**Authors:** Angana Mukherjee, Daniel P. Hollern, William A. Byrd, Tyeler S. Rayburn, Oluwasina G. Williams, Carrie Knight, Clayton C. Yates, Jacqueline. D. Jones

**Affiliations:** Department of Biological Sciences, Troy University, Troy, AL, United States; The Salk Institute, La Jolla, California, USA; Downstate-One Brooklyn Health, Brooklyn, NY, USA; Department of Pathology, The University of Alabama at Birmingham, Birmingham, AL, USA; Sidney Kimmel Comprehensive Cancer Center, Johns Hopkins School of Medicine, Baltimore, MD, USA; Department of Chemistry, Physics, and Engineering, Troy University, Troy, AL, United States

## Abstract

Studies have shown that all men who die from prostate cancer exhibit bone involvement, highlighting the clinical significance of bone metastases. Previously, we demonstrated that prostate cancer exhibits elevated expression of *FOXI3;* a transcription factor critical to bone development. However, its functional role in prostate cancer progression remains unexplored. In this study, we explored FOXI3 expression in human prostate cancer tissues and cell lines. Immunohistochemical analysis revealed a significant association between FOXI3 expression, tumor pathology, and increasing tumor grade. Analysis of publicly available gene expression data from prostate cancer patients showed that that *FOXI3* is markedly elevated in bone metastases and strongly correlates with *FGF8*, suggesting a potential bone-specific regulatory interaction between these factors. Consistent with this, treatment of bone-derived prostate cancer cells with FGF8 increased *FOXI3* RNA expression, whereas no such effect was observed in a brain-metastatic cell line. Further, we demonstrated that FGF8–FOXI3 axis regulates bone-derived prostate cancer cell migration and proliferation in a *FOXI3* dependent manner. Together, our findings demonstrate a pro-metastatic role of *FOXI3* in prostate cancer progression.

## INTRODUCTION

Prostate cancer is the most commonly diagnosed invasive solid tumor in men worldwide, representing approximately 7.3% of all cancers, with an estimated 375,000 deaths reported in 2022^1^. Even though only 5-7% of prostate cancer patients exhibit metastasis to distant sites, 60% of these patients die^2^. In particular, the death rate is 100% when prostate cancer metastasizes to bone^3^. Therefore, it is critical to understand the mechanisms and potential targets involved in prostate cancer metastasis and colonization of the bone.

Although the exact mechanisms underlying the sequential process of prostate cancer metastasis to bone are unclear, studies have shown that the abundance of growth factors in the bone microenvironment primes the cancer cells to proliferate and flourish^4^. The bone marrow microenvironment consists of a heterogeneous population of hematopoietic stem cells, osteoblasts, osteoclasts, bone stromal cells, bone transcription factors, and growth factors such as TGFβ, FGFs, IGFs, and BMPs^5–8^. Not surprisingly, many of these factors exhibit overlapping roles in both bone remodelling during early development and solid tumor progression^9–15^. Notably, the forkhead family transcription factor *FOXI3* has been reported to play a crucial role during early bone development^16–19^.

In previous work, we highlighted the role of *FOXI3* in various developmental processes and noted many of them also influence tumor progression^20^. We observed elevated *FOXI3* mRNA in a limited number of TCGA breast cancer bone metastasis samples and found that *FOXI3* mRNA expression increased with tumor stage in TCGA prostate cancer samples^20^. These data suggested that *FOXI3* may have important roles in tumor progression and bone metastasis. In this study, we expanded our analysis of prostate cancer patients and combined computational studies and functional studies in bone-derived prostate cancer cells, showing that *FOXI3* mRNA mediates metastatic phenotypes and demonstrating a potential mechanistic linkage to a key upstream growth factor, FGF8.

## MATERIALS AND METHODS

### Cell Culture, Antibodies and Reagents

Human prostate cancer cell lines DU-145, PC-3, and C4-2B were a gift from Dr. Laurie McCauley (University of Michigan, Ann Arbor) or purchased from ATCC. Cells were grown in DMEM with 10% FBS (Gibco) and 1% penicillin/streptomycin (Invitrogen) at 37 °C in 5% CO and were routinely confirmed to be mycoplasma-free.^21^.

### Immunohistochemistry

Prostate cancer tissue microarrays (TMAs) were obtained from TissueArray.Com LLC and UB Biolab (TMA# PRO811a and Pro162-02). Immunohistochemistry (IHC) was performed using the ABC kit (Vector Labs) and anti-FOXI3 antibody (Abcam#ab81824) as previously described^22^. Stained samples were blindly scored for cytoplasmic and nuclear FOXI3 intensity (0–4+) and the percentage of cells at each intensity was quantified using NIS Elements (Nikon).

### Quantitative RT-PCR

Total RNA was extracted using TRIzol (Invitrogen). cDNA was synthesized using universal RT-PCR kit (Applied Biosystems). *FOXI3* expression was measured using probes (Applied Biosystems, Hs03645828_s1; cat#4351372). Fold change was calculated and gene expression was quantified relative to *HRPT1* mRNA.

### RNA Interference Experiments

Cells were transfected 24 hours after plating with *FOXI3* specific or control siRNA (Origene) using Lipofectamine 2000 (Invitrogen). Post 24 hours of transfection, FOXI3 knockdown was confirmed by qRT-PCR, and only samples with >75% knockdown were used for experiments.

### Trypan Blue Assay

Trypan blue (Life Technologies) was used to determine cell viability. Cells were treated with varying FGF8 concentrations (0, 10, or 100 ng/mL) for 24 hours, trypsinized, mixed 1:1 with Trypan blue, and live cells were counted using a Countess II automated cell counter (Thermo Fisher Scientific).

### Migration Assay

siFOXI3 or siControl-treated, or untreated cells were seeded in 6-well plates, grown to 70% - 80% confluence, and starved overnight in Opti-MEM medium. A scratch was introduced in the monolayer^23^ followed by treatment with FGF8 (0, 10, or 100 ng/mL) in dialyzed media. Images were captured at 0 and 24 hours, and migration was quantified as the percentage reduction in the scratch area relative to baseline.

### Invasion Assays

Cancer cell invasiveness was determined using 24-well Transwell inserts (8 µm pores; BD Biosciences) following the manufacturer’s protocol. Matrigel-coated inserts were seeded with 6 × 10 cells and treated with 100 ng/mL FGF8 in serum-free media with 10% FBS in the lower chamber as a chemoattractant. After 24 hours, non-invasive cells were removed, and invading cells on the underside were fixed in 100% methanol, stained with 0.1% crystal violet, and counted in three random 10× fields.

### Statistical and Genomics Analysis

Statistical analyses were performed using GraphPad Prism v5.0 (GraphPad, La Jolla, CA). ANOVA and independent Student’s *t*-test were used to determine statistical differences between experimental and control values. *P* values <0.05 were considered statistically significant. Gene expression and copy number analyses were performed on published and publicly available molecular data featuring patient primary and metastatic samples^24,25^. Processed, normalized, gene expression and copy number data accessed on the cBIOPortal and analyzed using the portal’s built in plotting features^26,27^.

## RESULTS

### Increased FOXI3 expression is associated with high-grade prostate tumors

To evaluate FOXI3 protein expression and its association with prostate tumor grade, we performed IHC on TMAs comprising samples from 163 patients with varying prostate tissue pathology (Fig.1A and Supple.TableS1). Normal prostate tissues displayed no FOXI3 staining, while prostatic intraepithelial neoplasia (PIN) exhibited FOXI3 expression in the basal cell layer of ducts, suggesting early upregulation during tumorigenesis (<u>Figs.1A and 1B</u>). A comparison of FOXI3 expression across 91 patient tumor samples revealed significantly higher FOXI3 levels in grade 3–4 tumors compared with grade 1–2 tumors (<u>Figs.1A, 1C and Supple.TableS1</u>). In a separate analysis, we stratified 32 patient samples based on Gleason grade—a widely used clinical metric of prostate cancer aggressiveness. Tumors with Gleason grades 3 to 5 exhibited significantly higher FOXI3 expression than those with Gleason grades 1 or 2 (Figs. 1A, 1D, and <u>Supple.TableS1</u>). Collectively, these results suggest that elevated FOXI3 expression correlates with higher tumor grade and may serve as a marker of aggressive prostate cancer.

**Figure 1.**
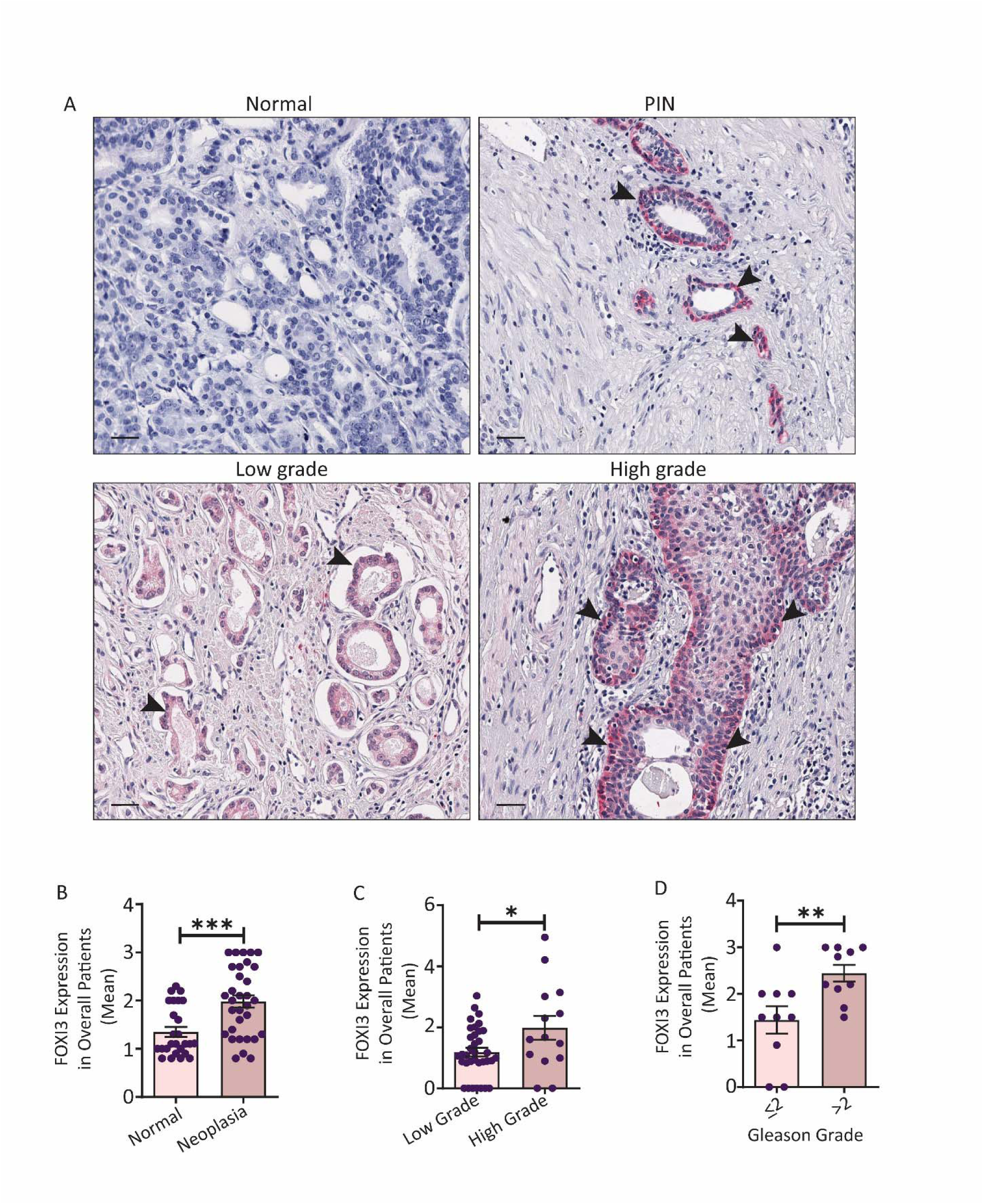
*FOXI3* expression in human prostate cancer specimens. **(A)** Representative images show FOXI3 expression from IHC studies of 163 prostate cancer tissues. Arrows indicate positive cells. Total magnification is 10X. PIN is Prostatic intraepithelial neoplasia. **(B-D)** FOXI3 staining intensity was quantified computationally for each specimen and plotted according to pathology type, tumor grade and Gleason grade (*P=0.0191, **P=0.0097, ***P=0.0004)

### FOXI3 expression is elevated with prostate bone metastasis and FGF8 expression

Building on the IHC data demonstrating elevated FOXI3 protein expression in high-grade prostate cancer, we hypothesized that FOXI3 expression may also correlate with metastatic progression. To test this, we analyzed gene expression data from two published studies featuring distant prostate cancer metastases^24,25^. While primary tumors showed a broad range of *FOXI3* mRNA levels, metastatic lesions generally exhibited higher expression, with bone metastases showing the highest *FOXI3* levels (Fig.2A and Supple.Figs.S1A and S1B). Notably, both primary and metastatic samples with the highest *FOXI3* expression often exhibited copy number gains at the FOXI3 locus, which were more common in bone metastases than in primary tumors (Fig.2A). Together, these data suggest that FOXI3 may contribute to bone-specific metastatic progression in prostate cancer and that copy number amplification could be a potential mechanism driving its overexpression in these lesions.

**Figure 2.**
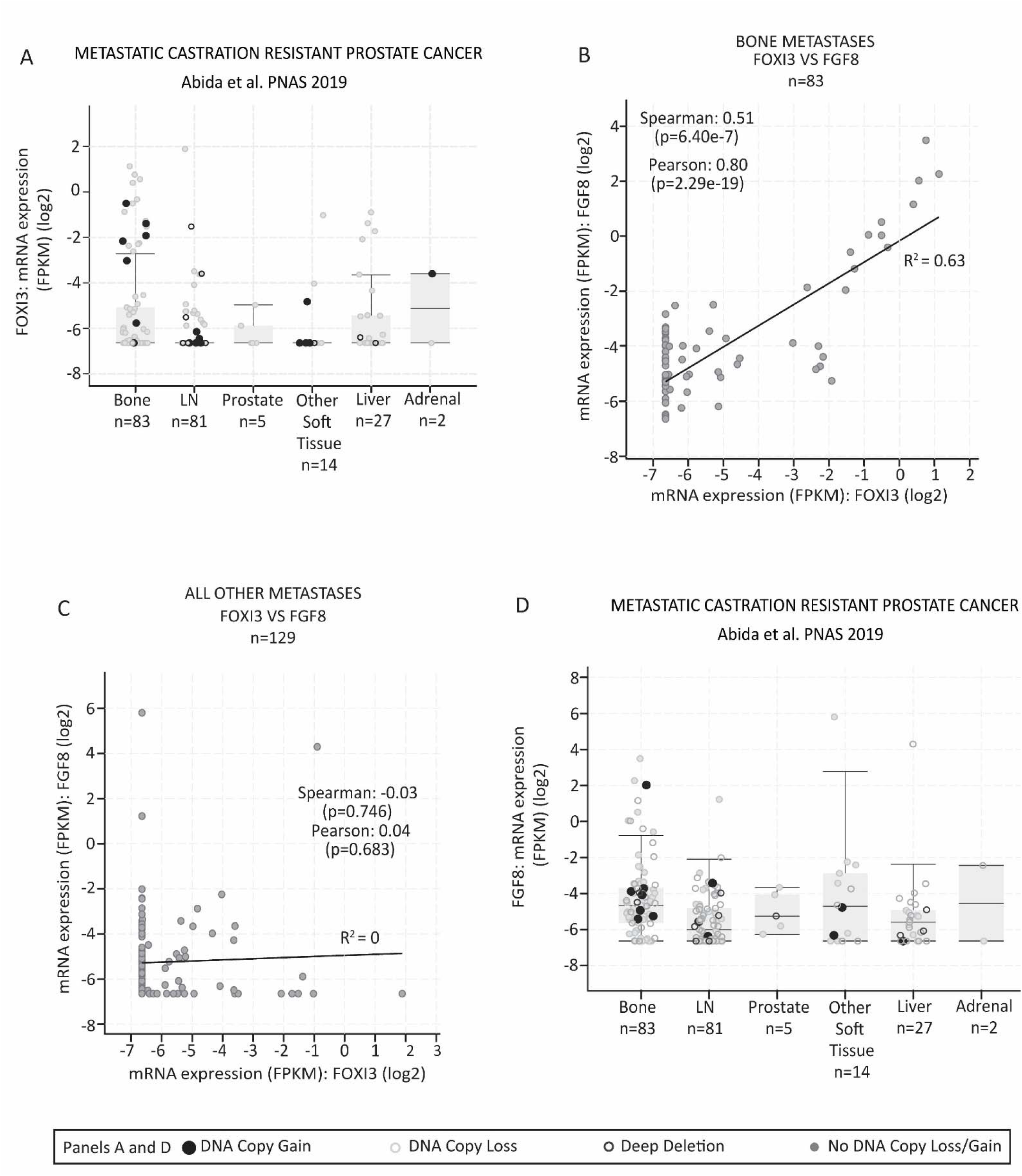
*FOXI3* expression and correlation with *FGF8* in prostate cancer metastasis. Gene expression and copy number analyses were performed using primary and metastatic patient samples from published and publicly available dataset (Abida et al., 2019)^25^. **(A)** Processed, normalized, *FOXI3* gene expression and copy number across metastatic sites was graphed to demonstrate FOXI3 expression patterns. **(B)** Spearman and Pearson correlation analysis show a significant correlation between *FOXI3* and *FGF8* in bone metastasis.. **(C)** No significant correlation between *FOXI3* and *FGF8* was observed in non-bone metastatic sites. **(D)** *FGF8* mRNA expression in prostate cancer metastasis, including bone.

Because FOXI3 acts as a transcription factor, we next analyzed patient datasets to identify genes whose expression strongly covaries with *FOXI3* expression, aiming to uncover potential upstream regulators that may serve as therapeutic targets for modulating *FOXI3* activity. Our analysis revealed a significant correlation between *FOXI3* and ***FGF8*** expression in tumors with bone metastases (Fig.2B and Supple.Fig.S1C); but not in other metastatic sites (Fig.2C). Moreover, similar to FOXI3, bone metastasis samples with high FGF8 expression also exhibited increased FGF8 copy number gains (Fig.2D). Together, these analyses suggest a potential functional link between FOXI3 and FGF8 that may be particularly relevant for prostate cancer progression in the bone microenvironment.

### FOXI3 is overexpressed in prostate cancer bone metastatic cells and regulated by FGF8

To identify suitable models for functional studies, we compared *FOXI3* expression in prostate cancer cell lines from bone metastases (PC3 and C4-2B) with brain-metastatic DU-145 cells^28,29^. qRT-PCR analysis showed that *FOXI3* expression was significantly upregulated in the bone-derived lines, 4-fold higher in PC-3 and 30-fold higher in C4-2B cells compared to brain-derived prostate cancer cell, DU-145 (Fig.3A), reflecting the increased *FOXI3* levels observed in patient bone metastases.

**Figure 3.**
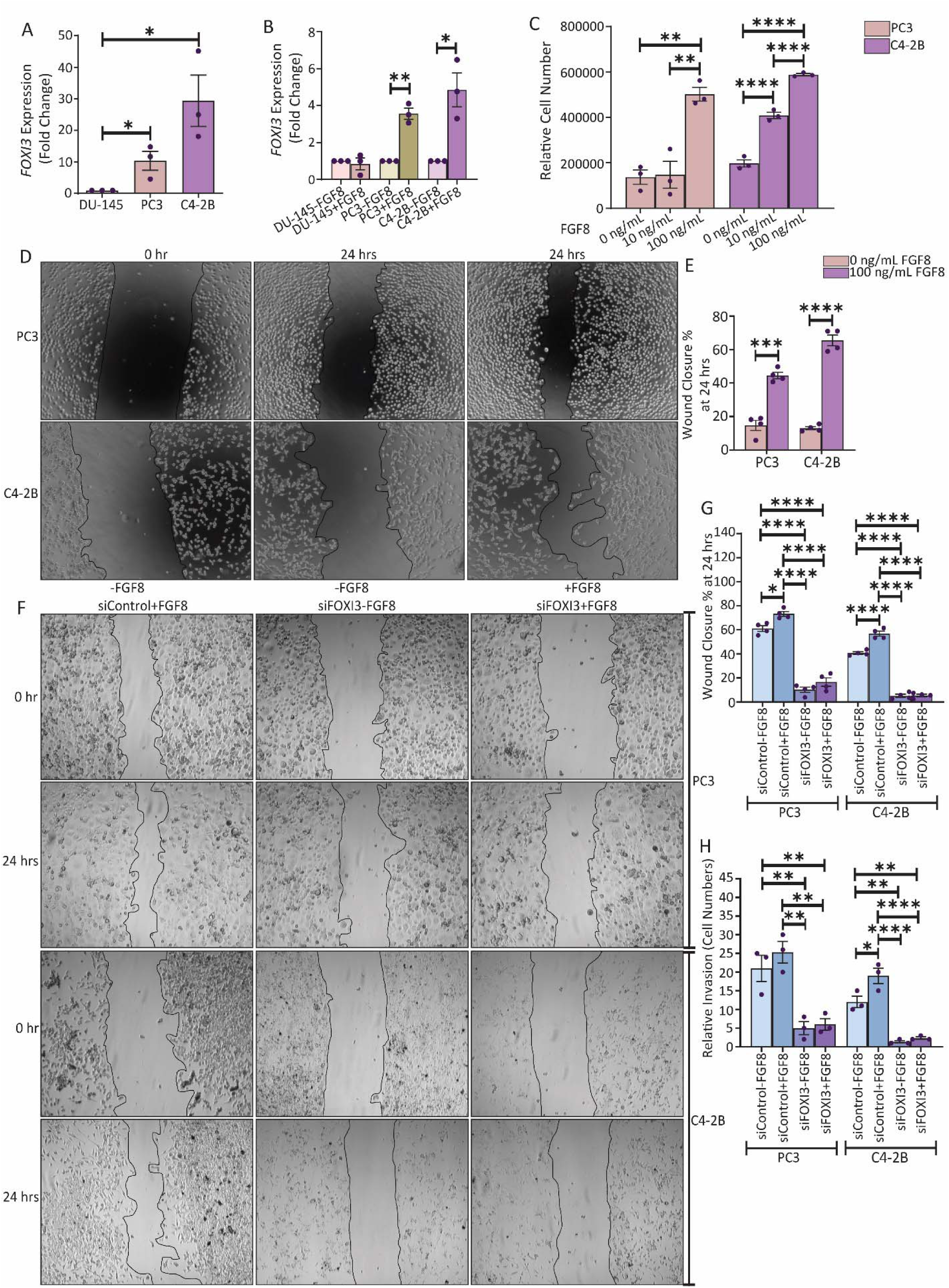
FGF8 elevates *FOXI3* expression and is required for FGF8-induced proliferation, migration, and invasion in bone-derived prostate cancer cell lines. **(A)** DU-145, PC3, and C4-2B prostate cancer cell lines were analyzed for *FOXI3* mRNA levels by quantitative RT-PCR (n=3 ± SE). HPRT1 was used as the endogenous control, and data was normalized to the brain tropic DU-145 cell line (*P= 0.0363, *P= 0.0250). **(B)** DU-145, PC3, and C4-2B prostate cancer cells were treated with 100 ng/mL FGF8, respectively, and *FOXI3* mRNA expression was analyzed after 24 hours by quantitative RT-PCR. GAPDH was used as the endogenous control. Data was normalized to the respective untreated cells line (n=3 ± SE) (*P= 0.0138, **P= 0.0011). **(C)** PC3 and C4-2B cells were treated with increasing concentrations of FGF8 (0ng/mL, 10 ng/mL, and 100 ng/mL) and after 24 hours proliferation was analyzed by trypan blue assay (n=3 ± SE) (PC: **P=0.0016, **P=0.0019; C4-2B: ****P<0.0001). **(D) and (E)** Post 24 hours of treatment with 100 ng/mL FGF8, a scratch-wound assay was performed on PC3 and C4-2B cells, respectively. Images were captured at a total magnification of 100X. Cell migration quantification was performed by measuring the distance between 3 random points within the wound edge in three replicate experiments. Data was normalized to 0 hours (n=4 ± SE) (PC: ***P=0.0002; C4=2B: ****P<0.0001). **(F) and (G)** 48 hours post transfection with siRNA against *FOXI3*, PC3, and C4-2B cells were treated with 100 ng/mL FGF8, and a scratch-wound assay was performed, respectively. Images were captured at a total magnification of 100X. Data was normalized to 0 hours (PC: *P=0.0119, ****P<0.0001; C4-2B: ****P<0.0001) (n=3 ± SE). **(H)** siFOXI3 transfected PC3 and C4-2B cells were plated on matrigel-coated filters, and an invasion assay was performed. Invasive cells were stained with crystal violet, and enumerated. (n=3 ± SE) (PC: **P=0.0056, **P=0.0082, **P=0.0013, **P=0.0018; C4-2B: *P=0.0138, **P=0.0011, **P=0.0021, ****P<0.0001)

Given the positive correlation between *FOXI3* and *FGF8* in prostate cancer bone metastasis (Fig.2B), and the known role of FGF8 in modulating FOXI3 during early development^16^, we hypothesized that FGF8 regulates FOXI3 activity in bone-metastatic prostate cancer. To test this, we treated prostate cancer cell lines with FGF8 and measured *FOXI3* expression. Consistent with patterns observed in patient metastasis data (<u>Figs.2B and 2D</u>), FGF8 selectively increased *FOXI3* levels in bone-derived PC3 and C4-2B cells, but had no effect on brain-derived DU-145 cells (Fig.3B). In addition, qRT-PCR analysis revealed a 30-fold and 80-fold increase in *FOXI3* mRNA expression in PC3 and C4-2B cells, respectively, compared to DU-145 following FGF8 treatment (<u>Supple.Fig.S2A</u>), exceeding the baseline *FOXI3* expression observed in untreated cells (Fig.3A). Together, these results indicate that FGF8 regulates *FOXI3* expression in a cell type–specific manner, particularly in bone-derived prostate cancer cells.

### FOXI3 and FGF8 promotes proliferation and migration in bone-metastatic prostate cancer cells

While our data showed that FGF8 regulates *FOXI3* expression in bone-derived prostate cancer cells, it remained unclear whether this interaction contributes to cancer cell progression. To address this, we treated PC3 and C4-2B cells with increasing concentrations of FGF8 (0, 10, and 100 ng/mL), and proliferation was measured after 24 hours. In PC3 cells, treatment with 100 ng/mL resulted in a significant increase in proliferation (Fig.3C), while C4-2B cells exhibited a dose-dependent response, with significant increases in proliferation observed at both 10 ng/mL and 100 ng/mL FGF8 (Fig.3C). To assess the effect of FGF8 on prostate cancer cell migration, we performed a scratch wound assay using PC3 and C4-2B cells. FGF8 treatment significantly increased the migration rate in both bone-derived lines PC3 and C4-2B (<u>Figs.3D and 3E</u>), whereas DU-145 cells, which express low levels of *FOXI3*, showed no significant change in the migration rate with FGF8 (<u>Supple.Figs.S2B and S2C</u>). These results suggest that FGF8 promotes a more aggressive, migratory phenotype specifically in *FOXI3*-high prostate cancer cells.

To determine whether these FGF8-driven phenotypes require FOXI3, we performed siRNA-mediated knockdown of *FOXI3* in PC3 and C4-2B cells (<u>Supple.Fig.S2D</u>). Reduced *FOXI3* expression markedly diminished FGF8-induced proliferation (<u>Supple.Fig.S2E</u>) and also significantly impaired migration and invasion in both cell lines, even without FGF8 stimulation (<u>Figs.3F–3H</u>). These findings show that *FOXI3* not only mediates FGF8-induced behaviors but also independently drives migratory and invasive phenotypes, underscoring the functional significance of the FOXI3-FGF8 axis in prostate cancer progression.

Together, results demonstrate that FGF8-mediated aggressive phenotypes are FOXI3-dependent, and that FOXI3 itself controls key processes required for metastatic progression. When combined with our computational analyses of patient datasets, these findings implicate FOXI3 as a key regulator of prostate cancer malignancy and a likely promoter of bone-tropic metastatic behavior.

## DISCUSSION

In this study, we integrated genomic datasets with immunohistochemical analysis of prostate tumor microarrays to investigate the role of FOXI3 in prostate cancer, with a focus on bone metastasis. Building on our earlier work implicating FOXI3 in cancer biology^20^, we found that FOXI3 expression increases with tumor grade, supporting its association with aggressive disease. Clinical datasets further revealed co-elevated expression of *FOXI3* and the bone-associated growth factor FGF8, specifically in bone metastases. Our functional assays demonstrated that this correlation has biological significance: FGF8 upregulates *FOXI3* in bone-derived prostate cancer cells, promoting increased proliferation and migration. To our knowledge, this is the first direct evidence of a functional role for FOXI3 in cancer, establishing it as a potential candidate effector in prostate cancer progression to bone.

Analysis of clinical datasets also revealed that a subset of prostate cancer patients exhibited copy number gains at the FOXI3 locus, with some also showing FGF8 amplifications. In both cases, these alterations were associated with increased gene expression and occurred more frequently in bone metastases than in primary tumors. Given that copy number changes are established drivers of metastatic progression^30,31^, our findings suggest that FOXI3 may similarly function as a driver in prostate cancer.

We also observed a significant correlation between FOXI3 and FGF8 in patients with prostate cancer bone metastases. This relationship was functionally validated by in vitro experiments, which showed that FGF8 promotes *FOXI3* expression, leading to increased proliferation and migration of prostate cancer cells derived from bone metastases. Notably, this correlation was not observed in other metastatic sites, suggesting that the FGF8–FOXI3 axis contributes specifically to bone tropism. These findings highlight the need for future in vivo studies to define FOXI3 downstream targets and to identify the cellular sources of FGF8, which may uncover therapeutic strategies to block bone metastasis or eradicate disseminated tumor cells. The association of FOXI3 with higher tumor grade, together with functional evidence showing its role in proliferation, migration, and invasion, supports the idea that tumor clones with elevated *FOXI3* and *FGF8*—likely resulting from genomic amplification —may have an enhanced capacity to metastasize to bone, underscoring the need for in vivo studies to define FOXI3 downstream targets and clarify the cellular sources of FGF8. Such work could inform therapeutic approaches aimed at blocking bone metastasis or eradicating disseminated tumor cells.

The observed copy number gains in FGF8 further suggests that tumor cells themselves may be a source of FGF8, consistent with earlier reports^32,33^. However, given the developmental roles of both FOXI3 and FGF8 in bone biology^34,35^, it is also plausible that the bone microenvironment, particularly osteoblasts, which are a known source of FGF8^36^, contributes to elevated FGF8 levels that potentially activates FOXI3 expression in tumor cells, thereby promoting proliferation and colonization. This aligns with the concept of osteomimicry, whereby prostate cancer cells adopt osteoblast-like programs to adapt to the bone niche^37,38^. Given their shared developmental roles, prostate cancer cells may exploit FGF8-driven FOXI3 activation to sustain survival and expansion in bone (Fig.4)

**Figure 4.**
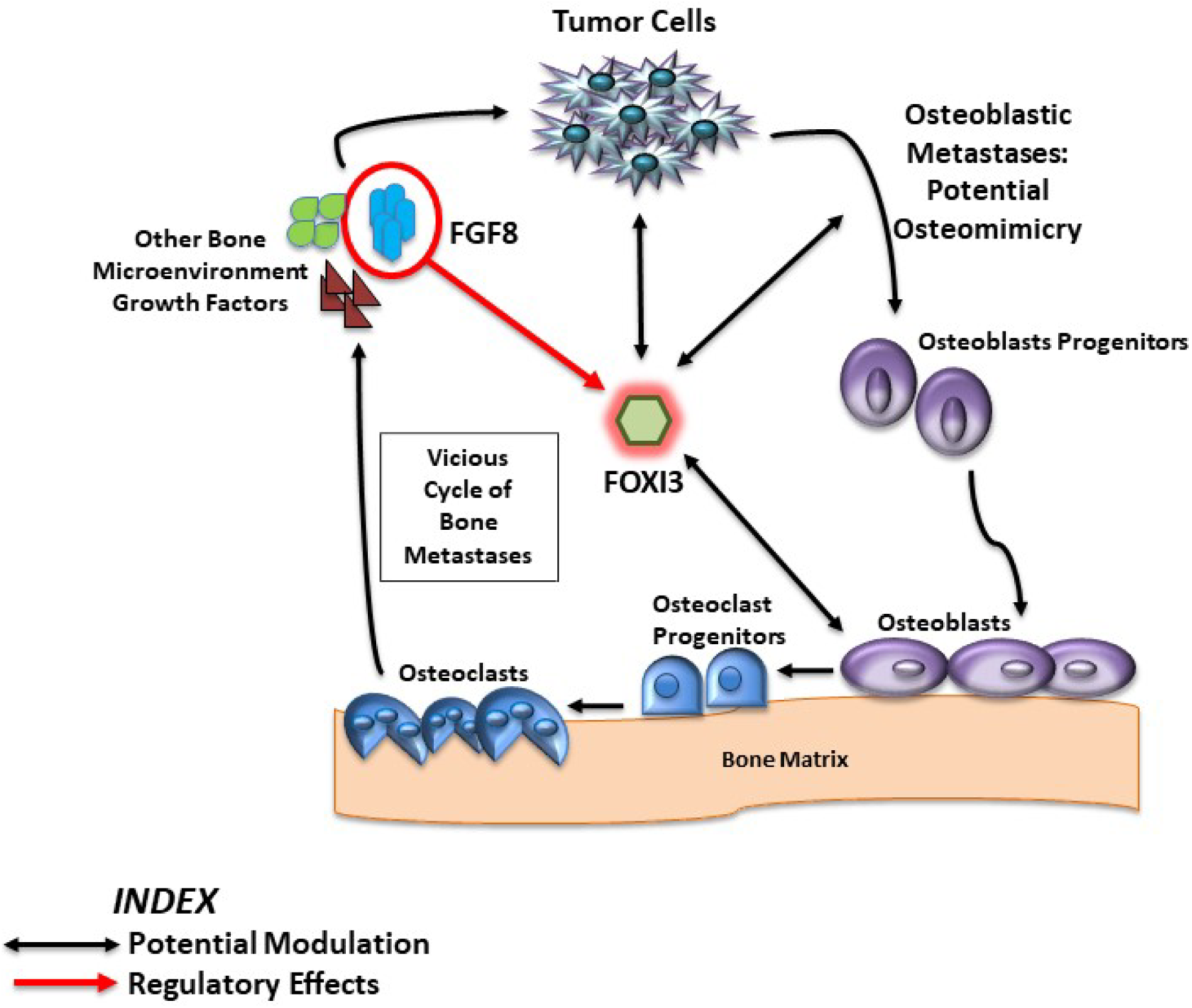
Proposed model depicting the role of *FOXI3* in prostate cancer bone metastasis. Metastatic prostate cancer cells may adopt osteoblast-like features, leading to enhanced FOXI3 activity that contributes to bone remodeling. Within this context, FOXI3 may be further modulated by the bone microenvironment factor FGF8, which amplifies tumor proliferation and colonization, thereby promoting bone metastasis.

In summary, this study provides the first patient-relevant evidence implicating FOXI3, in association with FGF8, in prostate cancer progression to bone. Supported by in vitro functional data, we show that *FOXI3* expression is closely linked to proliferation and migration of bone-derived prostate cancer cells. As a brief communication our findings open several avenues for future investigation, including identifying FOXI3 target genes, defining bone-derived regulators of FOXI3, and elucidating its role in osteomimicry. Together, these studies will be critical for determining whether FOXI3 and FGF8 can be leveraged as therapeutic targets in advanced prostate cancer with bone metastases.

## Supporting information

All Supplemental Figures

## Author contributions

A.M., J.D.J., and C.Y. conceived and designed the work. A.M., D.P.H and J.D.J wrote the manuscript. A.M. performed the IHC, RNA isolation, protein isolation, proliferation assays, migration assays, invasion assays, transient knockdown studies. D.P.H obtained and managed publicly available data; and performed the gene expression analysis. W.A.B., T.S.R, and O.G.W assisted with the cell culture, protein isolation and RNA isolation work. C.K. validated the IHC analysis. C.Y. provided reagents and cell culture support. All authors reviewed the manuscript.

