## Supplementary material for "FOXI3 Promotes Migration and Proliferation in Prostate Cancer Bone Metastases, Modulated by FGF8": All Supplemental Figures

**Supplementary Figures**

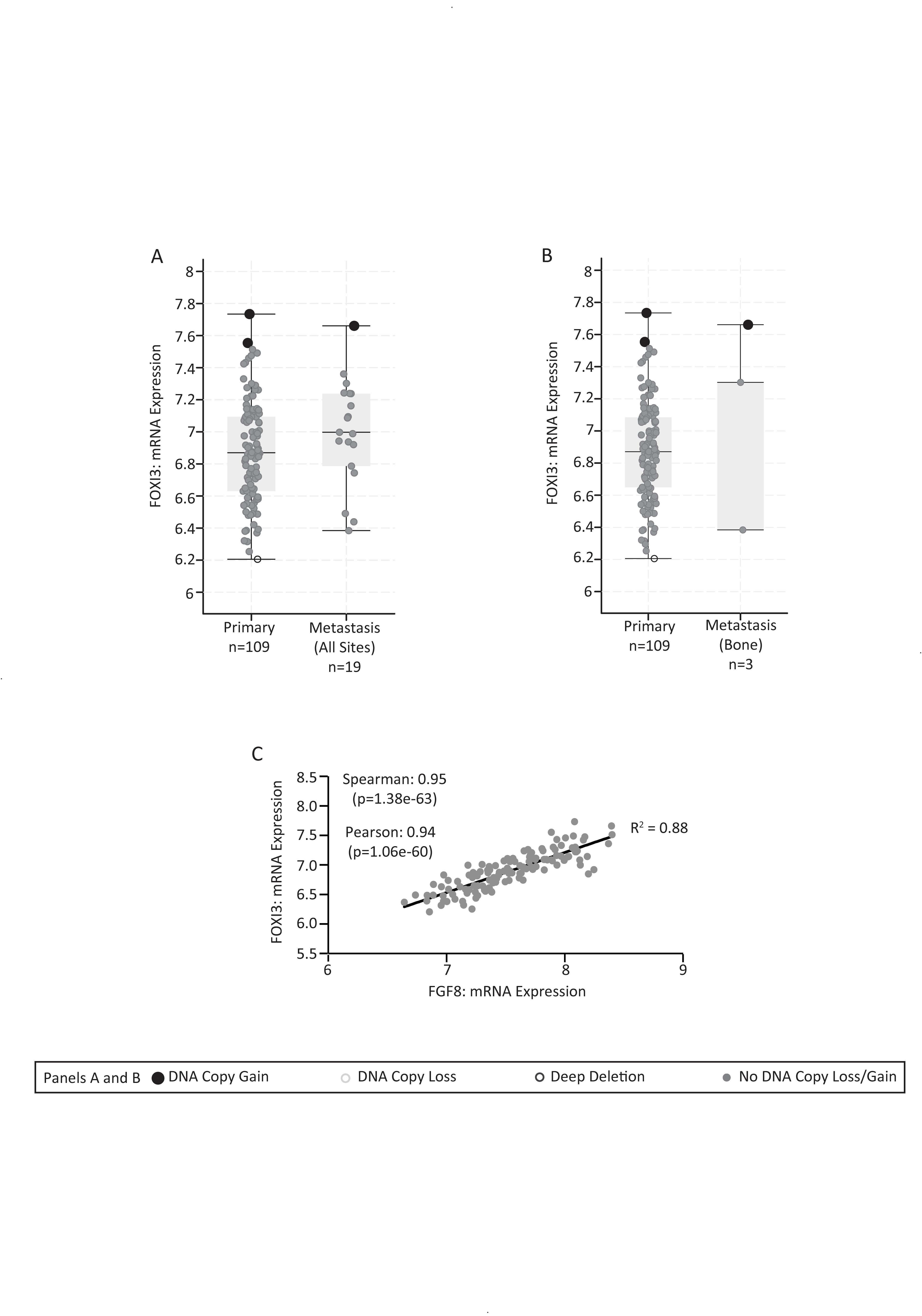

**Supplementary Figure S1.** ***FOXI3* and *FGF8* expression and correlation in prostate cancer patient samples.** Publicly available data (Taylor et al., 2010)^24^ was used to analyze *FOXI3* and *FGF8* gene expression and copy number. **(A)** *FOXI3* mRNA expression was graphed to demonstrate increased levels in metastatic sites compared to primary tumors. **(B)** Patients with bone metastases showed elevated *FOXI3* expression relative to primary tumors. **(C)** Spearman and Pearson correlation analysis revealed a significant positive correlation between *FOXI3* and *FGF8* in bone metastasis patient samples

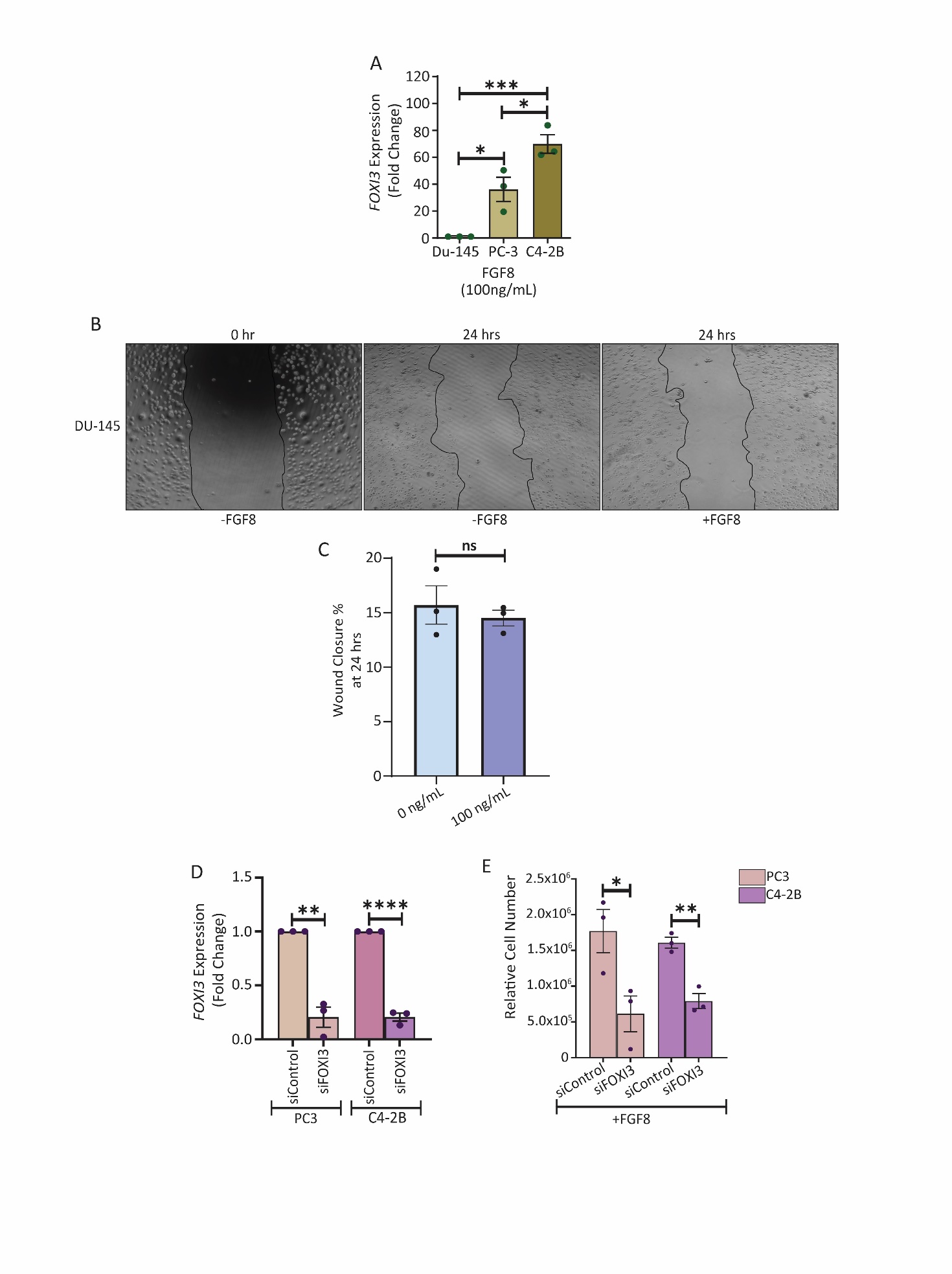

**Supplementary Figure S2.** **FGF8 regulation of *FOXI3* expression and functional effects on proliferation and migration in prostate cancer cell lines** **(A)** Post 24 hours of FGF8 treatment (100 ng/mL), *FOXI3* mRNA expression was analyzed in DU-145, PC3 and C4-2B prostate cancer cells by quantitative RT-PCR. HPRT1 was used as the endogenous control. Data was normalized to the brain metastatic DU-145 cell line (*n* = 3 ± SE) (^*^P=0.0173, ^*^P=0.0411, ^***^P<.001). **(B)** and **(C)** DU-145 cells were grouped into FGF8 treatment and non-treatment, and a scratch-wound assay was performed. Images were captured at a total magnification of 100X. Data was normalized to 0 hours (*n* = 3 ± SE). **(D)** Knockdown levels of *FOXI3* mRNA as determined by quantitative RT-PCR. HPRT1 as the endogenous control (*n* = 3 ± SE) (PC3: ^**^P=0.0010, C4-2B: ^****^P<0.0001). **(E)** siFOXI3 PC3 and C4-2B cells were treated with 100 ng/mL FGF8, and proliferation rate was assessed after 24 hours. (n=3 ± S.E) (PC: ^*^P=0.0419, C4-2B: ^**^P=0.0031).
